# A Multiscale Protein Abundance Structured Population Kinetic Model Systematically Explores the Design Space of Constitutive and Inducible CAR-T cells

**DOI:** 10.1101/2023.01.09.523295

**Authors:** Harshana Rajakaruna, Milie Desai, Jayajit Das

## Abstract

Engineered chimeric antigen receptor (CAR)-T cells are designed to bind to antigens overexpressed on the surface of tumor cells and induce tumor cell lysis. However, healthy cells can express these antigens at lower abundances and can get lysed by CAR-T cells. A wide variety of CAR-T cells have been designed that increase tumor cell elimination while decreasing destruction of healthy cells. However, given the cost and labor-intensive nature of such efforts, a systematic exploration of potential hypotheses becomes limited. To this end, we develop a framework (PASCAR) by combining multiscale population dynamic models and multi-objective optimization approaches with data obtained from published cytometry and cytotoxicity assays to systematically explore design space of constitutive and tunable CAR-T cells. We demonstrate PASCAR can quantitatively describe in vitro and in vivo results for constitutive and inducible CAR-T cells and can successfully predict experiments outside the training data. Our exploration of the CAR design space reveals that CAR affinities in an intermediate range of dissociation constants (K_D_) in constitutive and tunable CAR-T cells can dramatically decrease healthy cell lysis but sustain a high rate of tumor cell killing. In addition, our modeling provides guidance towards optimal tuning of CAR expressions in synNotch CAR T cells. The proposed framework can be extended for other CAR immune cells.

## Introduction

Chimeric antigen receptor (CAR)-T cells have been widely successful in treating hematologic cancers^1,2^. CAR-T cells are engineered to express CAR molecules which bind to antigens overexpressed in tumor cells and stimulate cytotoxic T cell responses. However, healthy cells can express such antigens at low copy numbers^3,4^ and can become targets of CAR-T cell cytotoxicity^5-7^. Such off-target destruction to healthy tissues is a major source of severe immune related adverse effects in patients undergoing CAR-T cell therapy^8,9^ and can become a major issue for solid tumors where viral organs are damaged by CAR-T cells. Therefore, generating an optimal CAR-T cell response that maximizes tumor cell elimination while minimizing off target destruction has been a longstanding pursuit of CAR-T cells therapies. A wide range of design strategies to engineer CAR-T cells have been proposed, including constitutive co-expression multiple CARs^10,11^ that sense several target antigens overexpressed by cancer cells or generation of inducible CAR expressions that tune copy numbers or abundances of CARs^6,12^ depending on the abundances of target antigens present on target cells. However, most of these design strategies are conceived intuitively, and thus limit systematic and wide explorations of the design space for CAR-T cells.

Optimal design strategies in economics or engineering often involve optimization of conflicting objectives where the best trade-offs between objectives are explored by subjecting computational/mathematical models to multiple objective optimization or Pareto optimization^13-15^. However, application of similar approaches for designing CAR-T cells face a major challenge due to the lack of experimentally validated and computationally efficient multiscale mathematical/computational models that can describe CAR-T cell responses against target cells. Though a large variety of population dynamic models based on ordinary differential equations (ODEs)^16^ have been developed to describe kinetics of populations of CAR-T cells interacting with cancer cells in vitro or in vivo, most of these models do not explicitly include CAR-ligand affinities, CAR abundances, co-receptors and their cognate ligand molecules, or biochemical signaling processes initiated upon engagement of CARs with cognate ligands. Since the design of CAR constructs often involves manipulation of the CAR affinities, CAR abundances, or T cell signaling processes, it is difficult to systematically explore the design space of CAR constructs with such existing models.

CAR abundances in single CAR-T cells can vary over 1000-fold in CAR-T cell populations^6^ which could play an important role in regulating the response of CAR-T cell populations against target cells. For example, in vitro cytotoxic assays of T cells with inducible CAR expressions showed that an increase in mean CAR abundances by less than 1.3-fold can produce over 3-fold increase in the rate of lysis of target cells in vitro. Furthermore, abundances of target antigens (e.g., CD19) on tumor cells in patients can also vary over 100-fold and lead to disparate (responders or non-responders) outcomes in patients undergoing CAR T cell therapy^17^. A handful of recent pharmacodynamic models^18,19^ incorporated CAR-ligand interactions by considering mean abundances of CARs and their cognate ligands, however, these models are unable to capture the variations of single cell CAR abundances or the T cell signaling kinetics. Detailed agent-based models incorporating details of receptor-ligand interactions, T cell signaling kinetics and cell metabolism have been developed to describe CAR-T cell responses^20^, however, quantitative validation of these models with experiments is challenging due to the presence of many difficult to calibrate model parameters and computationally intensive nature of the simulations.

We develop an experimentally validated <u>p</u>rotein <u>a</u>bundance <u>s</u>tructured population dynamic model for <u>CAR-</u> T cells (PASCAR) that integrates molecular receptor ligand interactions to single CAR-T cell signaling and activation to population kinetics of interacting CAR-T cells and target cells. PASCAR integrates CAR-ligand interactions and the ensuing signaling kinetics with a minimal but generalizable mechanistic modelling approach using ordinary differential equations (ODEs) where model parameters can be well estimated from routinely carried out experiments such as cytotoxicity assays and flow cytometry. We demonstrate PASCAR can quantitatively describe in vitro results for constitutive and inducible CAR-T cells and can successfully predict experiments outside the training data. Constitutive CAR-T cells constitutively express CAR moelcules whereas inducible CAR-T cells generate CAR expressions depending on its interaction with target antigens on target cells. PASCAR is then combined with a Pareto optimization that includes the trade-off between lysis of tumor and healthy cells to explore the design space of CAR constructs. Our investigations show CAR-ligand affinities with dissociation constants in the micromolar range can dramatically decrease healthy cell lysis but sustain a high rate of tumor cell killing. The proposed framework can be extended to model responses in other CAR immune cells^4,21,22^.

### Model Development

We developed a protein abundance structured model (PASCAR) for describing interacting populations of CAR-CD8+ T cells and target cells. In the model, individual CAR-T cells interact with single target cells where the interaction between a CAR-CD8+ T cell and a target cell is initiated by binding of the CAR with its cognate ligand such as HER-2. Once the CAR-ligand complex is formed, it goes through a series of chemical modifications such as phosphorylation of tyrosine residues in the CAR associated CD3ζ adaptors due to signaling reactions^23,24^. These modifications eventually lead to the release of lytic granules by the CAR-T cell which induce disintegration of the target cell membrane and eventual death of the target cell^25^. The signaling reactions can also induce proliferation of the CAR-CD8+ T cells^26,27^. There are a multitude of interconnected signaling processes involving many proteins, lipids, and ions that link the formation of the CAR-ligand complex to the lysis of target cells or to CAR-T cell proliferation^29^. In the model, we make simplifying assumptions to model these signaling reactions to relate the rate of target cell lysis and CAR-CD8+ T cell proliferation to the abundances of CAR-ligand signaling complexes. Since signaling reactions occur at faster time scales (∼minutes)^28^ compared to the time scales (∼ hours) of target cell lysis^29^ or CAR-T cell proliferation^30^, we assume that the rates of target cell lysis and T cell proliferation are influenced by the steady state values of the abundances of CAR-ligand complexes in our model. To simplify the notation, we denote CAR and its cognate ligand by R and H, respectively, and denote the single cell copy numbers or abundances of these proteins by the italicized versions of the symbols. In PASCAR, a CAR-T cell with *R* number of CARs interacts with a target cell with expressing *H* number of cognate ligands. R binds with H at a rate *k*_on_ to create the complex, R-H, which unbinds at a rate *k*_off_ (Fig. 1). The formation of the complex R-H induces a series of signaling reactions in the CAR-T cell that eventually leads to the activation of the CAR-T cell.

**Figure 1.**
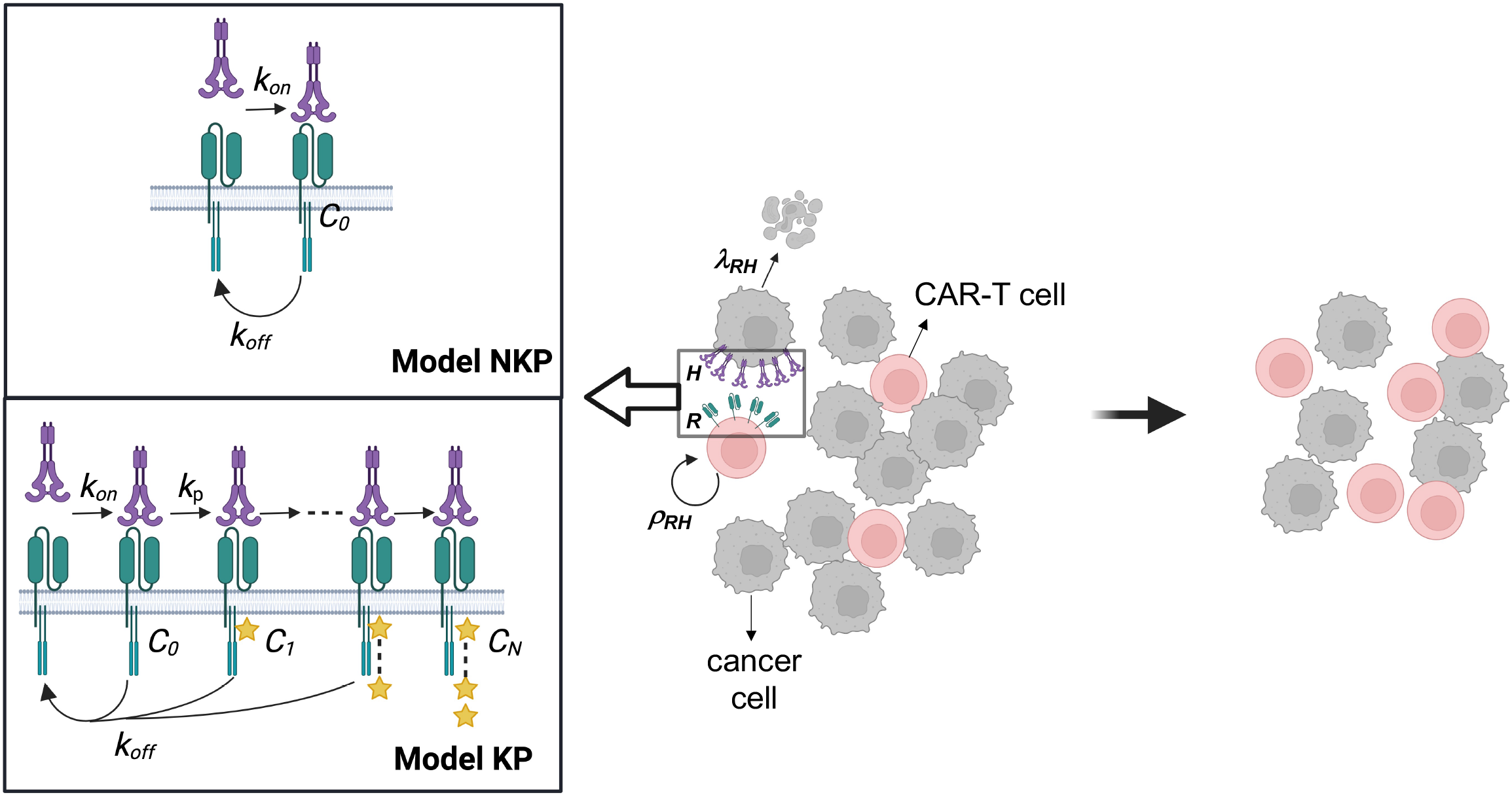
Multiscale PASCAR model. Single CAR-T cells interact with single target cells in the PASCAR model. The strength of CAR-T cell signaling depends on the abundance of the CAR-HER2 complex (C_0_) in Model NKP or on the abundance of an active complex (C_N_) formed due to *N* number of chemical modifications in the CAR-HER2 complex in Model KP. The abundances of C_0_ (Model NKP) or C_N_ (Model KP) depend on the abundances of CAR (*R*) and HER2 (*H*) in single CAR-T cell and the target cell, respectively. The rate of lysis (*λ*_*RH*_) of target cells due to cytotoxic response or the proliferation rate (ρ_RH_) of CAR-T cells depends on the abundances of C_0_ (Model NKP) or C_N_ (Model KP) in single CAR-T cells. The lysis and the proliferation rates are used to describe the kinetics of populations of CAR-T cells and target cells.

We propose two models, Model NKP and Model KP, to investigate different mechanisms of CAR-T cell activation due to CAR binding to cognate ligands. In Model NKP, CAR molecules in CAR-T cells interact with cognate ligands on target cells to form R-H complex and activation of a CAR-T cell is assumed to be proportional to the steady state abundance of the R-H complex (Fig. 1). In Model KP, we use an approximate model for CAR-T cell signal transduction based on McKeithan’s kinetic proofreading (KP) model^31^. In this model, the R-H complex formed in the CAR-T cell undergoes *N* number of modifications representing chemical modifications by downstream signaling reactions to create an active complex C_N_ (Fig. 1). Production of C_N_ leads to cytotoxic response and proliferation of CAR-T cells. The above series of reactions approximate signaling reactions in CAR-T cells initiated by the formation of CAR-ligand complex that produces activation (e.g., phosphorylation) of key signaling proteins^32,33^ such as Zap70 or PLCγ or SLP-76 crucial for CAR-T cell activation. The chemical modifications of the R-H complex are described by first order reactions (Fig. 1) with rate *k*_*p*_. At any chemically modified state of the complex, unbinding of the ligand leads to immediate reversion of all the modifications in the complex and the complex reverts to the native unbound ligand and receptor state. This step is known as the kinetic proofreading (KP) step which creates a sharp increase in the steady state number of C_N_ as *k*_*off*_ decreases. Thus, the KP model gives rise to a waiting time (τ_w_) that the receptor-ligand complex should last to generate productive signaling – receptor-ligand complexes with lifetimes (∼1/k_off_) larger than this waiting time (*i*.*e*., 1/k_off_ > τ_w_ ∝1/k_p_) generates active complexes C_N_ at greater abundances compared to short-lived (1/k_off_ ≪ τ_w_) receptor complexes. Below, we describe the kinetics of populations of interacting CAR-T cells and target cells for Model NKP and Model KP using ordinary differential equations (ODEs).

#### Model NKP

In this model, the rate of CAR-T cell proliferation or the rate at which a CAR-T cell lyses an interacting target cell is proportional to the steady state abundance of the R-H complex or C_0_. The abundance (*C*_0_) of the R-H complex formed at the steady state as a function of *R, H*, and the dissociation constant *K*_*D*_ = *k*_off_/*k*_on_ is given by,

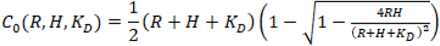

We assume that the rate of CAR-T cell proliferation ρ_*RH*_ and the rate of target cell lysis λ_*RH*_ are given by,

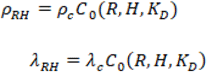

where, ρ_c_ and λ_c_ are the proportionality constants.

#### Model KP

The rates of CAR-T cell proliferation and target cell lysis are proportional to the steady state abundance of the end complex C_N_ which as a function of the steady state abundance of R-H or C_0_^(KP)^, k_p_, and k_off_ is given by,

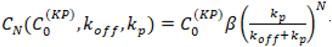

where, 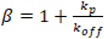 *C*_0_^(KP)^ is given by,

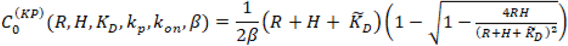

where,

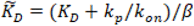

The proliferation and lysis rates in Model KP are given by,

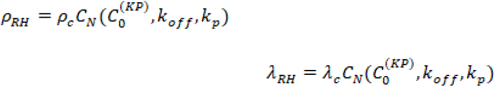

Now we set up kinetic equations for a population of target cells interacting with a population of CAR-T cells. The target cells can represent healthy or tumor cells. We consider a population of CAR-T cells where individual CAR-T cells express *R* copy number of CARs within a range [*R*_min_, *R*_max_]. The CAR-T cells interact with a population of target cells where any single target cell express *H* copy number of cognate ligands within a range [*H*_min_, *H*_max_]. The target cells replicate with a rate *r* and the target cell population decreases as they are lysed by interacting CAR-T cells. The CAR-T cells proliferate due to their interaction with the target cells, and the CAR-T cells die with a constant rate δ. The population kinetics can be described in terms of the number (*U*_*H*_) of target cells each carrying *H* copy number of cognate ligands and the number (*T*_*R*_) of CAR-T cells each carrying *R* copy number of CARs and as follows:

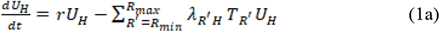

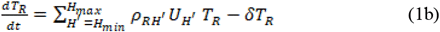

Given the initial condition {T_R_(0), U_H_(0)}, the above system of ODEs describe the composition or the structure of the populations of CAR T cells and target cells at any later time.

##### Parameter estimation

The affinity parameters (*k*_on_, *k*_off_) are usually measured in in vitro using experiments such as surface plasmon resonance^34^ or titrations using flow cytometry^35^. The single cell abundances of CARs and their cognate ligands are measured in quantitative flow cytometry experiments^6^ which can be used to estimate {T_R_(0), U_H_(0)}. The signaling parameter k_p_, the proportionality constants, λ_c_ and ρ_c_, can be estimated from cytotoxicity assays where CAR-T cells and target cells are co-cultured for different CAR-T cells and target cells ratios (or E:T ratios) and the fraction of lysed target cells is measured after few days (e.g., 3 days). We used data available from Hernandez-Lopez et al.^6^ for constitutive and synNotch-CAR-T cells to estimate parameters in our model. Further details are provided in the Materials and Methods section.

##### Pareto Optimization

There is a trade-off between maximizing lysis of cancer cells and minimizing lysis of healthy cells by CAR-T cells. For example, CAR-T cells that lyse target cells expressing a small number of cognate ligands can lyse cancer cells efficiently but can also produce large off target killing of healthy cells. We set up a multi-objective optimization problem to systematically explore the space of optimal parameter values in our PASCAR model and calculate the Pareto front, which represents the set of optimal parameter values where any objective cannot be optimized further without choosing less than optimal values for the other objectives. We set up a two objective optimization problem where the percentage of lysed healthy cells and the inverse of the percentage of the lysed cancer cells at a fixed time (e.g., 5 days) post co-incubation of the CAR-T cells and the target cells are minimized simultaneously. We consider a mixture of healthy and cancer cells where healthy and cancer cells express on average 10^4.5^ and 10^6.5^ HER2 molecules/cell^6^. The CAR-T cells are introduced at t=0 and interact with the target cells following the PASCAR model, and after a fixed time interval τ we evaluate the total numbers of lysed healthy and cancer cells. The parameters in the PASCAR model for constitutive and synNotch-CAR-T cells are varied to evaluate the Pareto fronts. Further details of the calculation are provided in the Materials and Methods section.

## Results

### 1. Model with kinetic proofreading captures lysis of target cells by constitutive CAR T cells

We evaluated the capability of Model NKP and Model KP to describe the response of constitutive CAR-T cells expressing anti-HER2 single chain antibody (scFv) against target cells expressing HER-2 in vitro. We fitted both the models to the percentage lysis data obtained from cytotoxicity assays of target cells co-cultured with constitutive CAR T cells and to abundances of CARs in the T cells in the co-culture assayed after 3 days^6^. We found that Model NKP is unable to describe the increase in percentage lysis (Fig. 2A) as the affinity of the CAR increases from high (K_D_=17.6 nM, k_off_ =9.0× 10^−5^ s^-1^) to low (K_D_=210 nM, k_off_ =6.8× 10^−4^ s^-1^), though the model fits the means and the variances of CAR abundances in CAR T cells at t=3 days reasonably well (Fig. S1). This is because the abundances of CAR-HER2 complexes formed in Model NKP for the high and low affinity CARs are roughly similar (Fig. 2B) which produces almost equal rates of lysis and proliferation for the CAR-T cells bearing the high and low affinity CARs. Next, we fitted Model KP to the same percentage lysis and CAR expression data which successfully captured the increase in the lysis by the CAR-T cells expressing high affinity CARs(Fig. 2C). The model also reasonably fitted the means and variances of the CARs expressed at t=3 days (Fig. 2D). The estimated parameters (Table 1) show that inclusion of kinetic proofreading in the signaling kinetics with and active complex formed at N≈7 steps can separate the CAR T cell responses between high and low affinity CARs for the same HER2 concentrations (Fig. 2D).

**Figure 2.**
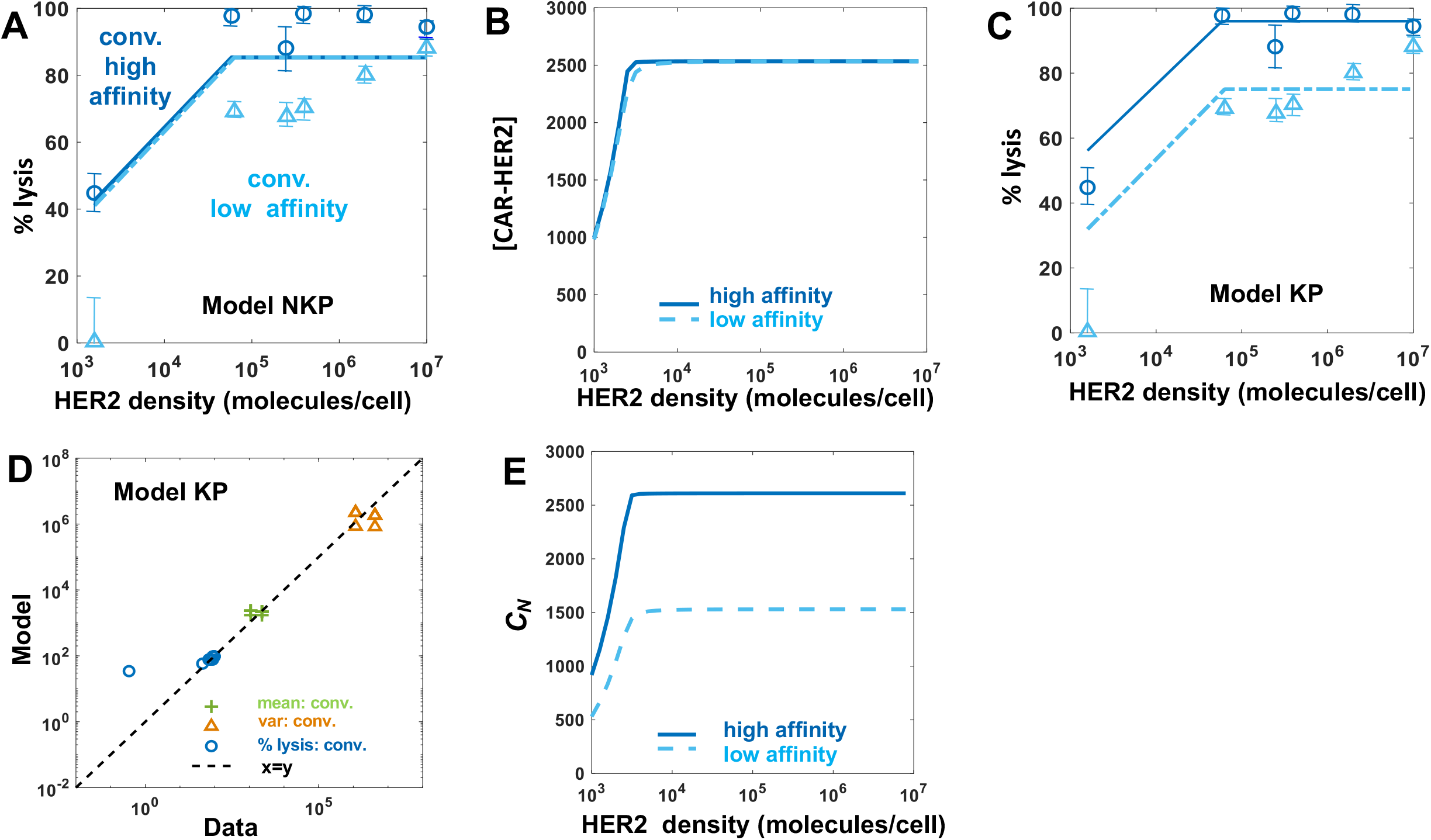
PASCAR modeling of cytotoxic and proliferation responses of constitutive CAR T cell against target cells. **(A)** Shows fits for Model NKP to the percentage lysis of target cells 3 days after 15,000 constitutive CAR T cells were incubated with 100,000 target cells. The data and the fits are shown for five different HER2 expressions with average HER2 abundances at 10^4.7^, 10^5.2^, 10^5.7^, 10^6.2^, 10^6.9^ molecules/cell. The constitutive CAR T cells express either high affinity (K_D_=17.6 nM, k_off_ =9.0× 10^−5^ s^-1^) or low affinity (K_D_=210 nM, k_off_ =6.8× 10^−4^ s^-1^) CARs. Variation of the CAR-ligand abundance (or C_0_) with the copy numbers of HER2 ligands expressed on target cells for the high and low affinity CARs in (A). **(C)** Shows fits for Model KP to the percentage lysis data described in (A). **(D)** Comparison of the model fits for Model KP and the experimental data for the means and the variances of CAR abundances. **(E)** Variation of the active CAR-ligand complex (or C_N_) with the copy numbers of HER2 ligands expressed on target cells for the high and low affinity CARs in (A). **(F)** Shows variations of the capacities of lysis (λ_RH_T_R_) and T cell proliferation (ρ_RH_U_R_) for different with time.

**Table 1:** List of estimated and fixed parameter values. The 95% confidence intervals are shown in squared brackets.

| Parameter | Unit | Constitutive +<br>synNotch | Constitutive |
| --- | --- | --- | --- |
| $\lambda_c$ | day<sup>-1</sup> | 2.09×10<sup>-8</sup><br>[1.78×10<sup>-8</sup>,<br>2.42×10<sup>-8</sup>] | 1.33×10<sup>-8</sup><br>[7.66×10<sup>-9</sup>,<br>2.05×10<sup>-8</sup>] |
| $k_p$ | s<sup>-1</sup> | 0.0074<br>[0.006, 0.009] | 0.0072<br>[0.0069,<br>0.0076] |
| N | number | 7.1961<br>[7.1734,<br>7.2187] | 6.9031<br>[6.7617,<br>7.0459] |
| $\rho_c$ | day<sup>-1</sup> | 5.31×10<sup>-9</sup><br>[3.76×10<sup>-9</sup>,<br>7.12×10<sup>-9</sup>] | 1.52×10<sup>-8</sup><br>[1.47×10<sup>-8</sup>,<br>1.58×10<sup>-8</sup>] |
| $\mu_c$<br>s.t. $\mu_R^{(consti.)}(0) = \mu_c$ | molecules/cell | 7.3589<br>[7.1719,<br>7.5483] | 7.2479<br>[6.9544,<br>7.5475] |
| $\sigma_R^{(consti.)}(0)$ | molecules<sup>2</sup>/cell | 0.4672<br>[0.4509,<br>0.4837] | 0.4841<br>[0.4748,<br>0.4935] |
| $\mu_s$<br>s.t.<br>$\mu_R^{(synN)}(0) = \mu_s \frac{\mu_H^{n_H}}{\mu_H^{n_H} + K_H^{n_H}}$ | molecules/cell | 6.5367<br>[6.2714,<br>6.8075] | - |
| $\sigma_R^{(synN)}(0)$ | molecules<sup>2</sup>/cell | 1.1257<br>[0.999,<br>1.2599] | - |
| $K_H$ | molecules/cell | 2.46×10<sup>5</sup><br>[2.46×10<sup>5</sup>,<br>2.46×10<sup>5</sup>] | - |
| $n_H$ | number | 3.8795<br>[3.7879,<br>3.9722] | - |
| **Fixed Parameters** |  |  |  |
| $k_{\text{off}}$<br>(high affinity CAR) | $\text{s}^{-1}$ | $9.0 \times 10^{-5}$ | $9.0 \times 10^{-5}$ |
| $k_{\text{on}}$<br>(high affinity CAR) | $\text{nM}^{-1} \text{s}^{-1}$ | $5.1136 \times 10^{-6}$ | $5.1136 \times 10^{-6}$ |
| $K_D$<br>(high affinity CAR) | $\text{nM}$ | 17.6 | 17.6 |
| $k_{\text{off}}$<br>(low affinity CAR) | $\text{s}^{-1}$ | $6.8 \times 10^{-4}$ | $6.8 \times 10^{-4}$ |
| $k_{\text{on}}$<br>(low affinity CAR) | $\text{nM}^{-1} \text{s}^{-1}$ | $3.2381 \times 10^{-6}$ | $3.2381 \times 10^{-6}$ |
| $K_D$<br>(low affinity CAR) | $\text{nM}$ | 210 | 210 |

Our PASCAR model allows us to analyze how CAR-T cell subpopulations expressing different CAR abundances respond to target cells. We used Model KP with best fit parameters (Table 1) to carry out the analysis. The ability of a CAR-T cell subpopulation expressing a specific CAR abundance to lyse target cells will depend on (i) the affinity (k_on_, k_off_) of CARs for HER2 and the strength of the ensuing signaling in the CAR-T cell, and (ii) the number of member CAR-T cells in the subpopulation. Therefore, a CAR-T cell subpopulation expressing higher CAR abundances but containing a small number of member cells could lyse target cells at a lower rate than a subpopulation expressing lower CAR abundances with a larger number of member cells. We quantified the rate of lysis of target cells expressing abundances of HER2 antigens of magnitude *H* by a CAR-T cell subpopulation expressing *R* number of CARs with *T*_*R*_ number of member cells by 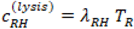. The kinetics of 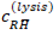 for target cells expressing mean *H* abundance (or 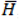) shows an increase with time resulting from the increase in *T*_*R*_ due to proliferation (Fig. 3A and Fig. S3). The CAR-T cell subpopulations with intermediate range of CAR abundances ∼(500 to 2500 molecules/cell) induce larger rate of lysis of target cells compared to the subpopulations with higher or lower CAR expressions (Fig. 3A). This is because the smaller number of CAR-T cells present in these subpopulations compared with that in subpopulations with intermediate CAR expressions. Moreover, the lower CAR abundances in the subpopulations with lower CAR expressions further reduce the cytotoxic response of the subpopulation. The rate of proliferation of CAR-T cell subpopulations as they interact with target cell subpopulations expressing specific abundances of HER2 antigens will depend on (i) the affinity (k_on_, k_off_) of CARs for HER2 and the strength of the ensuing signaling in the CAR-T cell, and (ii) the number of target cells expressing a particular abundance (H) of the HER2 antigens. The proliferation rate of a CAR-T cell subpopulation expressing abundances of magnitude *R* and containing T_R_ number of member cells as they interact with a target cell population of size *U*_*H*_ expressing abundance of magnitude *H* of HER antigens is quantified by 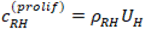. For a target cell subpopulation expressing mean abundance 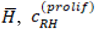, is given by 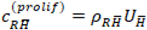. The proliferation rate of the CAR-T cells decreases with time as the number of target cells decreases over time due to the lysis of the target cells by the CAR-T cells (Fig. 3B and Fig. S2). The proliferation rates of CAR-T cell subpopulations increase with increasing abundances of CAR expressions (Fig. 3B and Fig. S2). This is because the signaling strength (∝ ρ_RH_) increases with increasing CAR abundances.

**Figure 3.**
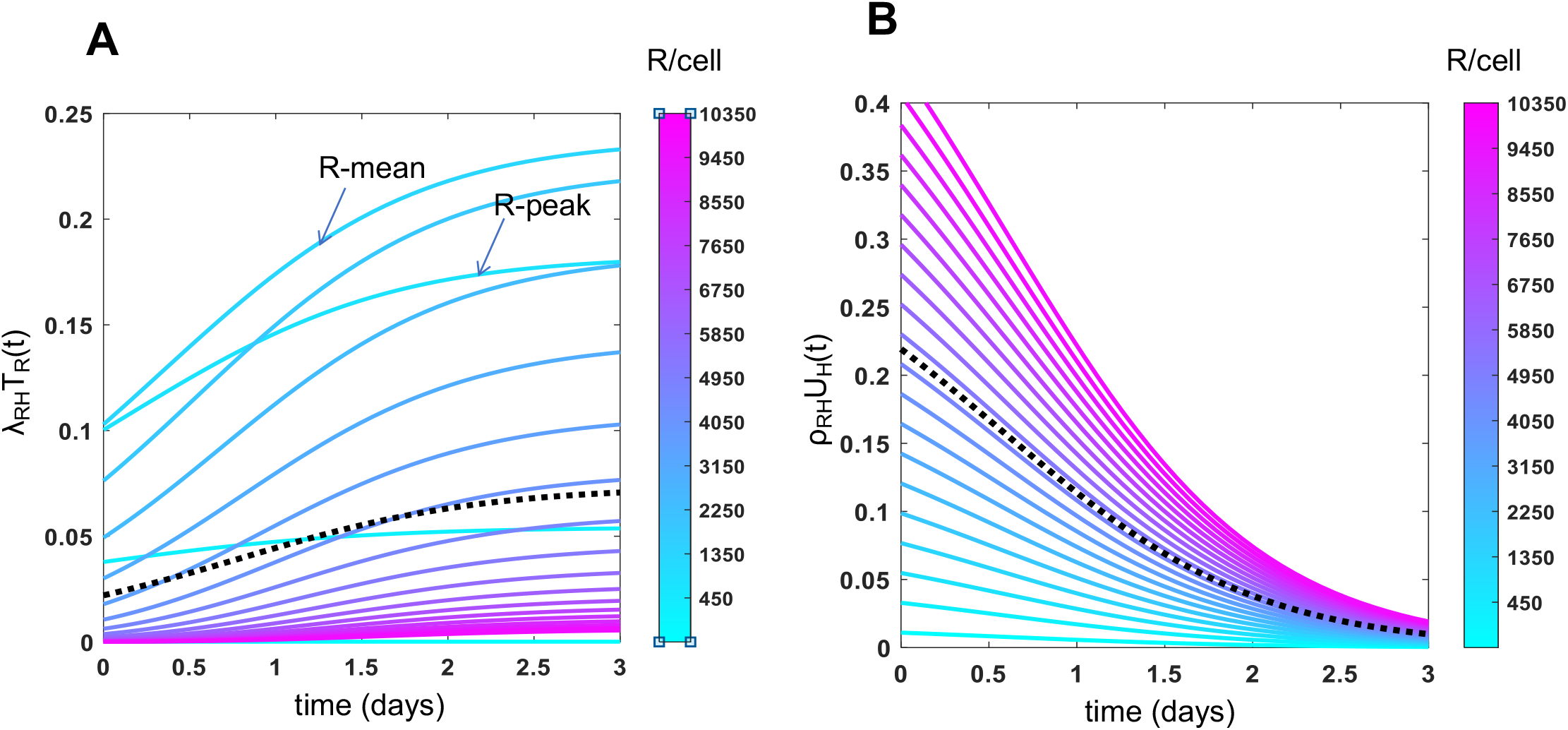
Constitutive CAR-T cell subpopulations expressing different CAR abundances show different cytotoxic and proliferative responses. **(A)** Shows lysis rates of target cells expressing mean HER2 abundances (=10^6.2^ molecules/cell) where the lysis is mediated by subpopulations of constitutive CAR-T cells expressing different CAR abundances. The lysis rates increase with time as the numbers of cells in the CAR-T cell subpopulations increase due to cell proliferation. The lysis rates for the CAR-T cell subpopulations corresponding to the mean CAR expression (denoted as R-mean) and to the mode of the CAR distribution (denoted as R-peak) at day 3 are marked. The dotted line shows the average of lysis rates across the CAR-T cell subpopulations. **(B)** Shows proliferation rates of CAR-T cells as they interact with target cells expressing mean HER2 abundances (=10^6.2^ molecules/cell).

### 2. Model with kinetic proofreading describes lysis of target cells by synNotch-CAR-T cells

We applied Model KP to describe cytotoxic response and proliferation of synNotch-CAR-T cells when the CAR-T cells were co-cultured with target cells in vitro. The mean CAR abundance of the synNotch-CAR-T cells in Model KP was assumed to be proportional to a Hill function of the mean HER2 abundance of the target cells (details in the Materials and Methods section). We reasoned that CAR-T cell signaling and activation in constitutive and synNotch-CAR-T cells should involve the same physiochemical processes, therefore, we fitted the percentage lysis data^6^ and the CAR expression data^6^ for constitutive and synNotch-CAR T cells simultaneously with Model KP. Model KP fits the percentage lysis and the means and variances of CAR and HER2 abundances reasonably well (Figs. 4A-B). The estimated parameters (Table 1) show a signaling cascade of N≈ 7 best fit the data. This result is consistent with recent experiments^36^ where clustering of LAT in T cell signaling is achieved in N=7.8± 1.1 kinetic proofreading reaction steps. The estimated value of the phosphorylation rate, k_p_ (≈ 0.007 s^-1^) is larger than the ligand unbinding rate k_off_ (≈ 10^−4^ s^-1^) indicating that the kinetic proofreading scheme is active in separating CAR-T cell responses across high and low affinity CARs. The estimated Hill function parameters (n_H_ ≈ 4, K_H_ ≈ 2 × 10^5^ molecules/cell) imply that CAR expressions are induced sharply as HER2 abundances increase past 2× 10^5^ molecules/cell giving rise to the almost binary cytotoxic response (off/on) against healthy cells (∼10^4.5^ HER2 molecules/cell) or tumor cells (>10^6.5^ HER2 molecules/cell). Next, we tested the ability of Model KP to predict synNotch-CAR-T cell response for experiments^6^ not included in training the model. We used Model KP to predict percentage lysis of target cells by synNotch-CAR-T cells at day 3 post incubation where the synNotch-CAR-T cells were co-incubated with different concentrations of target cells not included in model training. The model predictions captured the dependency of the cytotoxic response on the initial effector target ratio observed in experiments^6^ reasonably well (Fig. 4C). The analysis of the synNotch-CAR-T cell subpopulations to induce cytotoxicity showed that similar to constitutive CAR-T cells, synNotch-CAR-T cells with intermediate range of CAR abundances generated the larger response compared to the subsets with low and high CAR abundances (Fig. 4D and Fig. S3). The proliferation rates of synNotch-CAR-T cell subpopulations increase with increasing CAR expressions (Fig. 4E and Fig. S3) similar to that of constitutive CAR-T cells.

**Figure 4.**
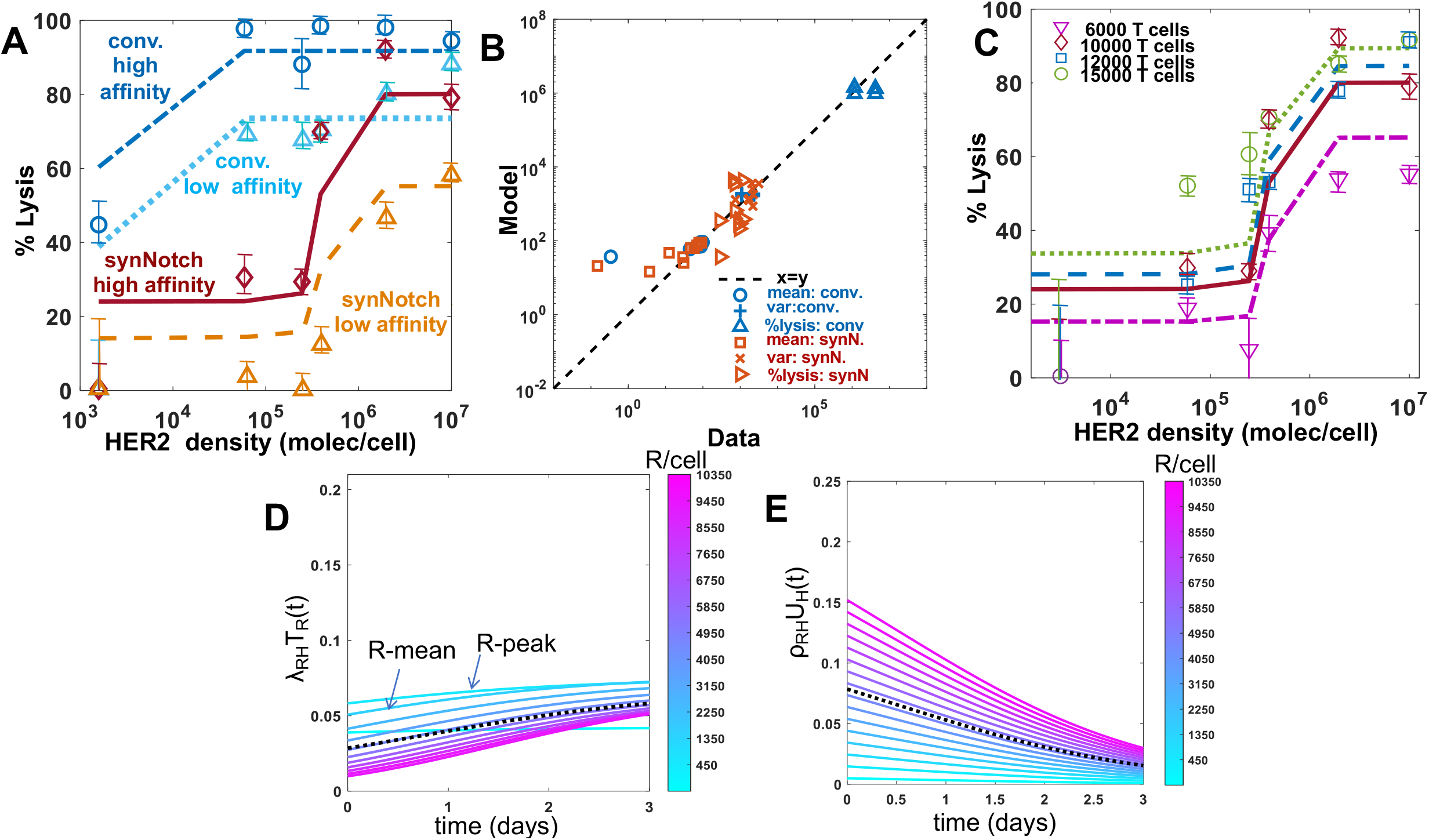
PASCAR modeling of cytotoxic and proliferation responses of constitutive and synNotch CAR-T cell against target cells. **(A)** Shows fits for Model KP to the percentage lysis of target cells at 3 days after 15,000 synNotch or constitutive CAR T cells were incubated with 100,000 target cells. The data and the fits are shown for five different HER2 expressions with average HER2 abundances at 10^4.7^, 10^5.2^, 10^5.7^, 10^6.2^, 10^6.9^ molecules/cell. The synNotch and constitutive CAR T cells express either high affinity (K_D_=17.6 nM, k_off_ =9.0× 10^−5^ s^-1^) or low affinity (K_D_=210 nM, k_off_ =6.8× 10^−4^ s^-1^) CARs. **(B)** Comparison of the model fits for Model KP and the experimental data for the means and the variances of CAR abundances at day 3 after co-incubation of synNotch or constitutive CAR T cells and target cells. **(C)** Shows PASCAR model predicted percentage lysis (solid and dashed lines) of target cells at 3 days after 6000, 10,000, and 12000 synNotch CAR-T cells were incubated with target cells. The predictions are in very good agreement with the data (shown using different symbols) obtained from Hernandez-Lopez et al. ^6^. The data for 15,000 synNotch CAR-T cells were used for training the model which is also shown as a reference. **(D)** Shows lysis rates of target cells expressing mean HER2 abundances (=10^6.2^ molecules/cell) where the lysis is mediated by subpopulations of synNotch CAR-T cells expressing different CAR abundances. **(E)** Shows proliferation rates of synNotch CAR-T cells as they interact with target cells expressing mean HER2 abundances (=10^6.2^ molecules/cell).

### 3. Optimal design of constitutive and inducible CAR-T cells

We performed Pareto optimization in the space of parameters that can be potentially manipulated in experiments. For constitutive CAR-T cells we carried out the analysis for CAR-ligand binding affinity parameters, k_on_ and k_off_, and, for synNotch-CAR-T cells, in addition to the CAR affinity parameters, the threshold (K_H_) and sharpness (n_H_) of CAR expression, were also considered. To evaluate the optimal parameters, we set up in silico cytotoxicity assays, where a mixture of 20,000 healthy and tumor target cells with 10,000 constitutive or synNotch-CAR-T cells were co-incubated in vitro (Fig. 5A). The CAR-T cells and the target cells interacted following Model KP set at the best fit parameter values (Table 1) and the percentages of healthy and tumor cells lysed after 5 days of culture were computed. When synNotch-CAR-T cells were present we assumed the CAR expressions in the synNotch-CAR-T cells are determined by the HER2 expressions of the tumor cells. The multi-objective optimization was performed in MATLAB where the competing objectives of percentage lysis of healthy cells and 1/(% lysis of tumor cells) were minimized simultaneously. The calculations of Pareto fronts showed that for constitutive CAR-T cells, decreasing k_off_ increases lysis of both healthy and tumor cells on a Pareto front at a fixed dissociation rate K_D_ = k_off_/k_on_. Decreasing k_off_ increases the lifetime of the CAR-HER2 complexes allowing for the complexes to last longer than the waiting time required for the signaling reactions to generate the active signaling complex required to activate the CAR-T cell. Thus, decreasing k_off_ leads to increase in the destruction of tumor and healthy target cells.

**Figure 5.**
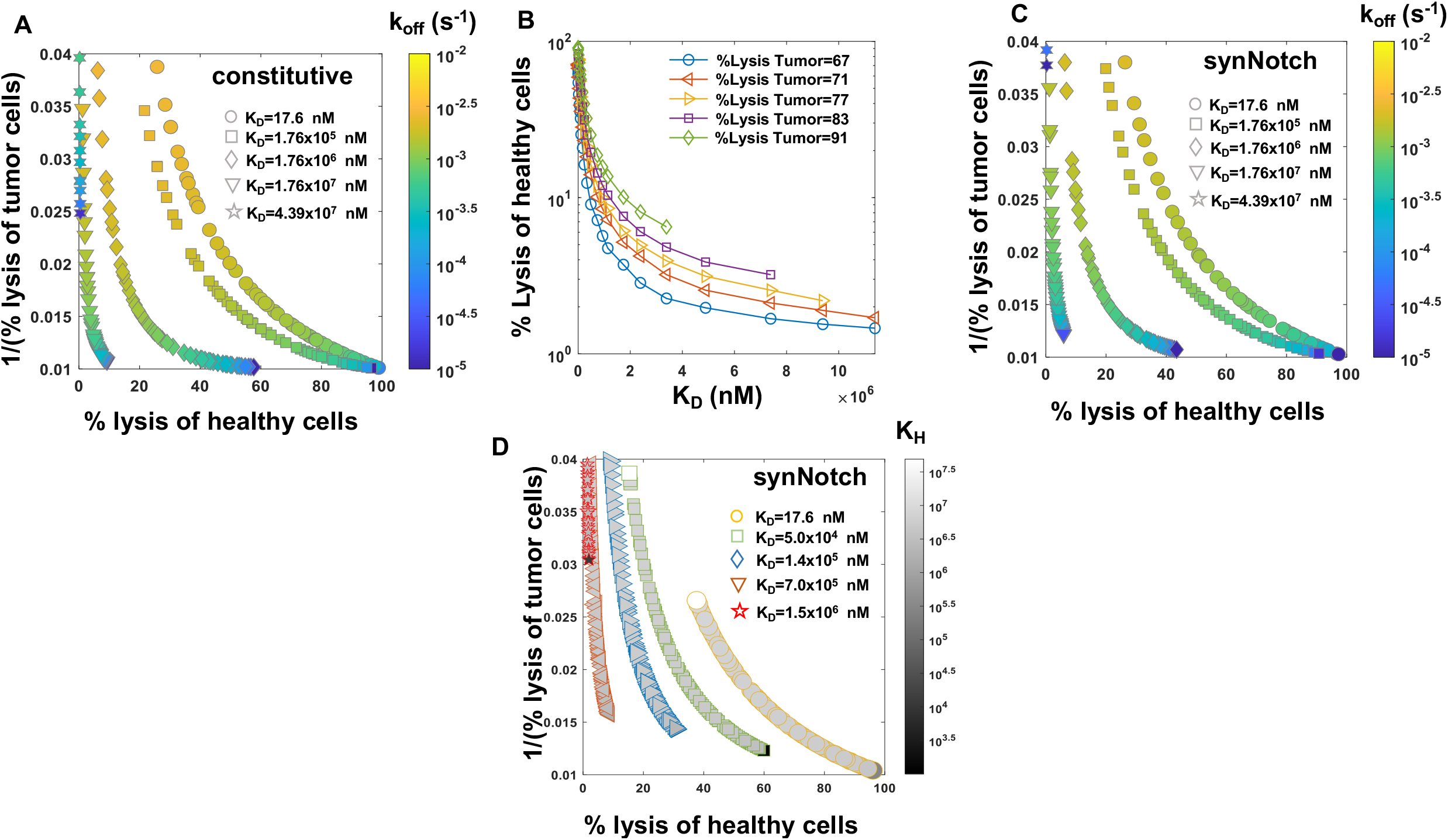
Pareto fronts revealing optimal responses of constitutive and synNotch CAR-T cells against tumor and healthy cells. **(A)** Shows Pareto fronts for constitutive CAR-T cells in the plane spanned by % lysis of healthy cells and 1/(% lysis of tumor cells). The Pareto fonts are calculated for different dissociation constants K_D_ = k_off_/k_on_ where for each K_D_ value, k_off_ and k_on_=k_off_/K_D_ are varied to obtain the corresponding front. The Pareto fronts are calculated 5 days after the CAR-T cells were incubated with a 1:1 mixture of tumor and healthy cells in silico (details in the main text). **(B)** Shows variations of % lysis of healthy cells with K_D_ for specific % lysis of tumor cells by constitutive CAR-T cells along the Pareto fronts. The % lysis of healthy cells decrease with increasing K_D_ until a certain value (the end points shown on the graph), increasing the K_D_ further decreases % lysis of tumor cells as well (not shown on the graph). **(C)** Shows Pareto fronts for synNotch CAR-T cells for different dissociation constants K_D_ = k_off_/k_on_ where for each K_D_ value, k_off_ and k_on_ =k_off_/K_D_ are varied to obtain the corresponding front. The other parameters K_H_ and n_H_ are fixed throughout at 2.42×10^5^ molecules/cell and 3.97, respectively. The in silico cytotoxic assay is set up the same way as in (A). **(D)** Shows Pareto fronts for synNotch CAR-T cells for different fixed K_D_ and k_off_ values at (3.8×10^−5^ s^-1^, 17.6nM), (1.0×10^−8^ s^-1^, 1.76×10^5^nM), (5.0×10^−9^ s^-1^, 1.76×10^6^ nM), (5.0×10^−9^ s^-1^, 1.76×10^7^nM) and (1.0×10^−9^ s^-1^, 4.39×10^7^ nM) where k_on_ =k_off_/K_D_. K_H_ and N are varied within [1×10^3^, 5×10^7^] molecules/cell and [1, 8] on each Pareto front. The symbol size is proportional to N and the shades filling the symbols are proportional to K_H_. The in silico cytotoxic assays are set up the same way as in (A).

However, Pareto fronts at different K_D_ values revealed that increasing K_D_ until a certain limit can increase the lysis of tumor cells while decreasing healthy cell lysis (Fig. 5A). For example, for a fixed % lysis of tumor cells (e.g., 66%), increasing K_D_ can decrease %lysis of healthy cells from 60% to 20% (Fig. 5A-B). However, decreasing K_D_ further starts decreasing lysis of tumor cells as well. This behavior can be explained by the dependency of the abundances of CAR-ligand complexes with K_D_. Since, the average abundances (∼ 5000 molecules/cell) of CARs are ∼6 and ∼600 times smaller than that of HER2 on tumor and healthy cells, respectively, most of the CARs in the CAR-T cells form complexes with HER2 antigens displayed by the healthy and the tumor cells for high affinity CARs (K_D_ ≪ 5000 molecules/cell). As, K_D_ is increased (K_D_ > 5000 molecules/cell), larger numbers of CAR-HER2 complexes are formed when CAR-T cells interact with tumor cells compared to the healthy cells due to the availability of 100 times more HER2 antigens on tumor cells. However, as K_D_ is increased further (K_D_ ≫ 5000 molecules/cell), the copy numbers of CAR-HER2 complexes decrease substantially for CAR-T cells interacting wth tumor and healthy cells as the majority of the CARs are unable to form complexes due to the weak affinity of the binding (Fig. S4). This results in inefficient lysis of healthy and tumor cells by CAR T cells. The above increase in discrimination between healthy and tumor cells with increasing K_D_ upto a certain limit is qualitatively consistent with experiments^37^ with CAR-T cells interacting with target cells expressing low (∼30,899 molecules/cell^37^) and high (∼ 628,265 molecules/cell^37^) EGFR abundances where the CAR was generated from high affinity cetuximab (K_D_=1.9 nM^38^) or low affinity nimotuzumab (K_D_=21 nM^38^). The experiments showed that the CAR-T cells with nimotuzumab produced a lower killing of target cells with low EGFR abundances compared to the CAR-T cells with cetuximab^37^. Both CAR-T cells produced similar killing of target cells with high EGFR abundances^37^.

Next, we carried out our analysis for synNotch-CAR-T cells. Similar to constitutive CAR-T cells, the Pareto fronts for synNotch-CAR T cells for fixed n_H_ and K_H_ values showed increased tumor- and decreased healthy-cell lysis when K_D_ decreased for a fixed unbinding rate k_off_ until a particular limit (Fig. 5C). For a fixed HER2 affinity (fixed k_on_ k_off_), increasing n_H_ and decreasing K_H_ increased lysis of tumor as well as healthy cells (Fig. 5D). This is because, increasing n_H_ and decreasing K_H_ produces higher CAR expressions in the synNotch-CAR T cells when they interact with tumor cells, and those CAR-T cells induce greater lysis of tumor cells. However, the same CAR-T cells become efficiently activated by healthy cells for higher affinity CARs (K_D_ ≪ 5000 molecules/cell, k_off_ ∼ 10^−5^ s^-1^) and thus induce increased lysis of healthy cells. Thus, an optimal design of these inducible CAR-T cells could be generation of CAR expressions with moderate affinities (K_D_ ∼ 1 μM, k_off_ ∼ 10^−4^ s^-1^) with higher values of n_H_ and lower values of K_H_.

## Discussion

We developed multiscale mechanistic model PASCAR that integrates processes across the scales of molecules of CARs and antigens to the populations of CAR-T- and target cells. This multiscale approach is particularly relevant for modeling response of heterogeneous populations of CAR-T cells expressing a wide range CAR abundances against target cells displaying antigen abundances that can vary over 1000 fold across target cells. We pursued an approximate or coarse-grained modeling approach where many microscopic details of CAR-T cell signaling and activation were incorporated implicitly in model parameters. This reduced approach allowed us to quantitatively explore the roles of CAR and HER2 (CAR antigen) abundances, CAR affinities, and the ensuing cell signaling in regulating CAR-T cell responses against healthy and tumor cells. The ability of this approach to describe the percentage lysis data and the proliferation of T cell subpopulations with different CAR abundances in cytotoxic assays, and to predict results outside model training shows the success of our approach in modeling anti-tumor responses of constitutive and inducible CAR T cells. We were also able to estimate model parameters reasonably well using flow cytometry measurements of CAR and HER2 expressions and percentage lysis data from cytotoxic assays. Therefore, PASCAR model combined with data from standard immunoassays can be used to make quantitative predictions for various experimental conditions investigating CAR-T cell responses in vitro.

Application of PASCAR to describe cytotoxic responses of constitutive CAR-T cells showed the relevance of kinetic proofreading in CAR-T cell signaling in discriminating response between high and low affinity CARs. Model NKP which did not contain any kinetic proofreading was unable to separate the cytotoxic responses mediated by high (K_D_=17.6 nM, k_off_=9.0×10^−5^ s^-1^) and low affinity (K_D_=210 nM, k_off_=6.8× 10^−4^ s^-1^) CARs. Including kinetic proofreading steps at early time signaling events in Model KP was able to separate the cytotoxic responses. There are several differences in signaling events induced by TCR and CAR stimulation, e.g., LAT is weakly phosphorylated in CAR-T cells^33^ whereas LAT is robustly phosphorylated in TCR signaling. Therefore, signaling proteins that could correspond to the active complex in Model KP could represent different signaling proteins in TCR and CAR signaling, e.g., experiments with light-gated immunoreceptors show the active complex corresponding the N≈7.8 in TCR signaling could represent LAT^36^, however, the active complex in Model KP for N=7 is likely to represent a different signaling protein.

We carried out a multi-objective optimization that minimizes destruction of healthy cells while maximizing the elimination of tumor cells when a select set of model parameters that can be manipulated in experiments were allowed to vary. The calculation of Pareto fronts in our multi-objective optimization showed that intermediate values of CAR affinities led to an increase in tumor cell killing while decreasing healthy cell killing. This finding is supported in previous experiments which showed reduction of CAR affinity reduced healthy cell killing but increased tumor cell killing^37^. For inducible CAR-T cells, the increase in the sharpness of CAR upregulation in conjunction with intermediate range of CAR affinities can produce a desirable amount of tumor cell and healthy cell killing.

An exciting extension of PASCAR would be to describe CAR-T cell response in vivo. CAR T cells undergo a maturation process over a longer duration (∼weeks) than that considered here to give rise to exhausted^39^ and memory phenotypes^34,40^. Cytokines and chemokines in the tumor microenvironment contribute to this maturation process^41,42^. These processes have been modelled for T cells^43^ and CAR-T cells^44^ using mechanistic or data-driven models which could be potentially incorporated in the PASCAR model.

### Limitations

The PASCAR model approximates signaling using a minimal model which might not be able to describe engineered CAR-adaptors that manipulates the number of ITAMs^17^, or engineered T cells where specific signaling proteins such as RasGTPase activating protein^45^ are knocked out. However, some of these changes could be potentially included in our minimal signaling network by implementing signaling models that have described similar effects in other contexts^46^. The kinetics of the induction and the decay^12^ of synNotch CARs are not considered in the current model which could be relevant to describe responses of synNotch CAR-T cells as they transit from microenvironments rich in tumor cells to healthy cells in vivo. An extension of the current model to include different compartments with explicit kinetics of synNotch CAR abundances could potentially address this issue.

## Materials and Methods

### Solution of the ODEs

We set up a rectangular lattice for the abundances *R* and *H* with lattice constants Δ_R_ and Δ_H_, respectively. *T*_*R*_ and *U*_*H*_ in the ODEs in Eq. (1) denote the numbers of CAR-T- and tumor cells with *R* and *H* abundances between *R* to *R*+Δ_*R*_ and *H* to *H*+Δ_*H*_, respectively. The ranges of R, Δ_*R*_, H, and Δ_*H*_, are chosen based on the ranges ([*R*_min_, *R*_max_] and [*H*_min_, *H*_max_]) and the distributions of R and H measured in flow cytometry experiments in ref. ^6^. Given the parameters, k_on_, k_off_, k_p_, λ_c_, ρ_c_, and the initial distributions of *R* and *H*, the nonlinear system of ODEs is solved numerically in MATLAB using the *ode45* function with Runge-Kutta 4 numerical method. The units of *R* and *H* in the ODEs are in (# of molecules)/cell.

### Unit conversion of kinetic rates

The rates k_on_, k_off_, and K_D_ are usually provided in literature in units of (nM)^-1^s^-1^ (or (μM)^-1^s^-1^), s^-1^, and nM (or μM), respectively. We convert k_on_ and K_D_ rates into units of (# of molecules)^-1^ s^-1^ and (#of molecules) to use in the ODEs where the units for CAR and HER2 abundances are given by (# of molecules)/cell. The unit conversion is carried out as follows. The nM unit is changed to (# of molecules)/(μm)^3^ using 1 nM = 600 × 10^−3^ (# of molecules)/(μm)^3^ =0.6 (# of molecules)/(μm)^3^. We assume CARs and HER2 molecules form complexes when these molecules are separated by a distance (*d*) of 2 nm^47^ or smaller, and the CAR and HER2 molecules interact in the immunological synapse formed at the interface of the interacting CAR-T cell and the target cell. The area (*A*) of the synapse region is taken as 1/2 times the area of a T cell^48^ = ½ × 4π (7/2)^2^ μm^2^. The number of CAR and HER2 molecules present in the area *A* is assumed to be half of the total numbers of these molecules present in individual cells. The parameters K_D_ and k_on_ in the units of (#of molecules) and (# of molecules)^-1^ s^-1^ are obtained by the following relations: K_D_ [# of molecules] = K_D_ [(# of molecules)/(μm)^3^]/(*Ad*)[(μm^3^)] and k_on_ [(# of molecules)^-1^s^-1^] = k_on_ [(# of molecules)^-1^s^-1^/(μm)^3^]/(*Ad*) [(μm^3^)].

### Parameter estimation

We estimated model parameters k_p_, λ_c_, ρ_c_, and the mean (μ_R_(0)) and the variance (σ_R_(0)) of CAR abundances at t=0 for constitutive and synNotch CAR T cells by fitting percentage lysis data and means and variances of CAR-T cells obtained at day 3 post incubation with target cells in cytotoxic assays. For synNotch-CAR-T cells, additional parameters describing upregulation of CARs due to binding of synthetic Notch receptors with HER2 ligands on target cells were estimated. We minimized the cost function^49^ below using *levenberg-marquardt* algorithm in MATLAB.

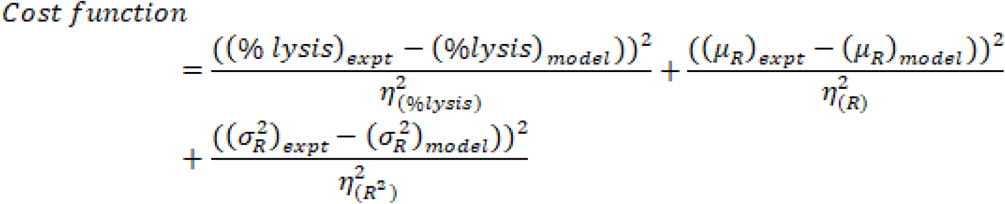

where, μ_R_ and σ_R_ denote the mean and the standard deviation of the CAR abundances at day 3 post incubation. 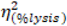 denotes the variance in the %lysis in the experiments which were calculated from ref ^6^ (Fig. 2A in that reference). 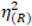 and 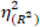 denote the variance and the fourth cumulant in CAR expressions, respectively, which were calculated from ref. ^6^. The confidence intervals were estimated by the *nlparci* function in Matlab using the residuals and covariance matrices given by the *nlinfit* in Matlab.

We provide specific details of parameter estimation for constitutive CAR and synNotch-CAR-T cells below. 1. *<u>Constitutive CAR-T cells</u>*: The data are obtained from Hernandez-Lopez et al.^6^ where human CD8+ T cells were engineered to express CARs constitutively that bind human epidermal growth factor receptor 2 (HER2) with high (K_D_ =17.6 nM) and low (K_D_ = 210 nM) affinities. In their experiments, human leukemia K562 cell lines were engineered to express five different average concentrations (10^4.2^, 10^5.2^, 10^5.7^, 10^6.2^, 10^6.7^ molecules/cell) of HER2 molecules which were used as target cells in cytotoxic assays. We fitted (*nlinfit* function in MATLAB) flow cytometry data (Fig.1C in ref. ^6^) with log-normal distributions to estimate means and variances of HER2 in target cells used in our ODE models. Similarly, the distributions of CAR abundances at day 3 post co-incubation were obtained by fitting the flow cytometry data (Fig. S1C in ref. ^6^) for constitutive CARs with log-normal distributions. We assumed the same CAR distributions for high and low affinity CARs. Means and variances for the CAR abundances at day 3 in the co-culture experiments were calculated from the estimated log-normal distributions. 2. *<u>synNotch-CAR-T cells</u>*: Hernandez-Lopez et al. ^6^ developed tunable CAR-T cells by engineering a synthetic notch (synNotch) receptor in CD8+ T cells. The synNotch receptor binds with HER2 on target cells with low affinity (210 nM) and acts as a high-density-antigen filter for inducing CAR expressions in the T cells. The generation of CARs by the synNotch circuit was modeled implicitly. We assume that the changes in the CAR expression in the syn-Notch CAR-T cells occur at a faster time scale than that of target cell lysis and T cell proliferation. Thus, the mean CAR abundance (*μ*_*R*_(0)) in synNotch-CAR T cells at the start of a co-culture experiment is assumed to be a Hill function of the mean HER2 abundance (μ_H_) of the target cells given by, 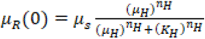. n_H_ is the Hill co-efficient and for n_H_≥2, μ_R_(0) changes in a switch like all or none fashion as μ_H_ increases beyond the threshold K_H_. The parameters μ_s_, n_H_, and K_H_ were estimated by fitting the percentage lysis (Fig. 2A in ref. ^6^) and the CAR expression data (Fig.S1C in ref. ^6^) obtained at 3 days after co-incubating target cells and CAR-T cells in cytotoxic assays. The estimated distributions of HER2 in target cells described for constitutive CAR-T cells were used for modeling experiments with synNotch-CAR-T cells as well.

#### Pareto optimization

A target cell population of 20,000 cells composed of a 1:1 mixture of healthy and tumor cells was taken at t=0. The healthy and tumor cells expressed HER2 molecules following a linear superposition of two lognormal distributions where mean values of the HER2 molecules were set to 10^4.5^ molecules/cell and 10^6.9^ molecules/cell for the healthy and tumor cells, respectively. The standard deviations of each lognormal distribution were set to 0.3 to avoid any substantial overlap between the distributions (Fig. S5). In our in silico cytotoxicity assays we considered the above 20,000 target cells were mixed with 10,000 constitutive or synNotch CAR-T cells at t=0 expressing a high affinity CAR (K_D_=17.6 nM, k_off_=9× 10^−5^ s^-1^). The distribution of CARs for the constitutive CAR-T cells were constructed following lognormal distributions with the parameters estimated in Table 1. For the synNotch CAR-T cells we assumed all the 10,000 CAR-T cells expressed CARs following a lognormal distribution with a mean 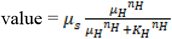 where *μ*_*H*_ =10^6.9^ molecules/cell. The values of n_H_ and K_H_ were set to the values estimated in Table 1 or varied for the Pareto front calculations. We used *gamultiobj* routine in Matlab to compute the Pareto front by optimizing two conflicting objective functions: f_1_ = % lysis of healthy cells at day 5, and f_2_ = 1/(% lysis of the tumor cells) at day 5. The Pareto fronts were calculated for fixed K_D_ values where k_off_ were varied. These calculations (Fig. 5C) for the synNotch CAR-T cells fixed the values of n_H_ and K_H_ at the values shown in Table 1. For these calculations we varied f_1_ and f_2_ as functions of k_off_ within ranges [1.0 ×10^−5^, 1.0 ×10^−2^] s^-1^ such that k_on_ =k_off_/K_D_ for different values of K_D_ fixed at (17.6, 1.76 ×10^5^, 1.76×10^6^, 1.76×10^7^, 4.39×10^7^) nM with all the other parameters fixed at the estimated values given in Table 1 for constitutive and synNotch CAR-T cells.

Another set of Pareto fronts were calculated for synNotch CAR-T cells (Fig. 5D) where n_H_ and K_H_ values were varied and all the other parameters including K_D_ and k_off_ were fixed. In this case f_1_ and f_2_ varied with n_H_ and K_H_ within ranges [1, 8] and [1×10^3^, 5×10^7^], respectively, with k_off_ and K_D_ values fixed at (3.8×10^−5^ s^-1^, 17.6nM), (1.0×10^−8^ s^-1^, 1.76×10^5^nM), (5.0×10^−9^ s^-1^, 1.76×10^6^ nM), (5.0×10^−9^ s^-1^, 1.76×10^7^nM) and (1.0×10^−9^ s^-1^, 4.39×10^7^ nM).

## Data and Code Availability

The MATLAB codes and the data used are available at the GitHub link https://github.com/Harshana4532/CART_Project_Matlab_03Jan2023.

## Acknowledgement

This work is supported by the Research Institute at the Nationwide Children’s Hospital. We thank members of the Das lab for discussions and feedback.

## Supplimentary Figure Captions

**Figure S1.**
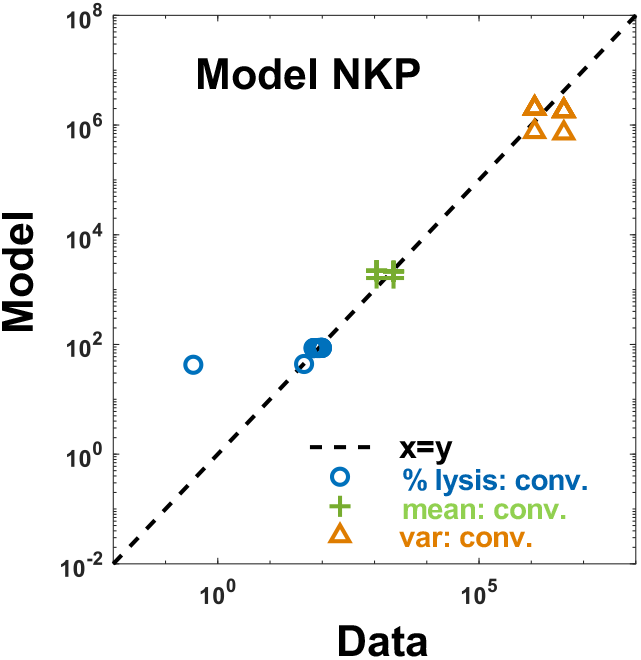
Fits to CAR expression and percentage lysis data in constitutive CAR-T cells for Model NKP. Shows model fits for Model NKP to the % lysis data and the means and the variances of CAR abundances at day 3 post co-incubation.

**Figure S2.**
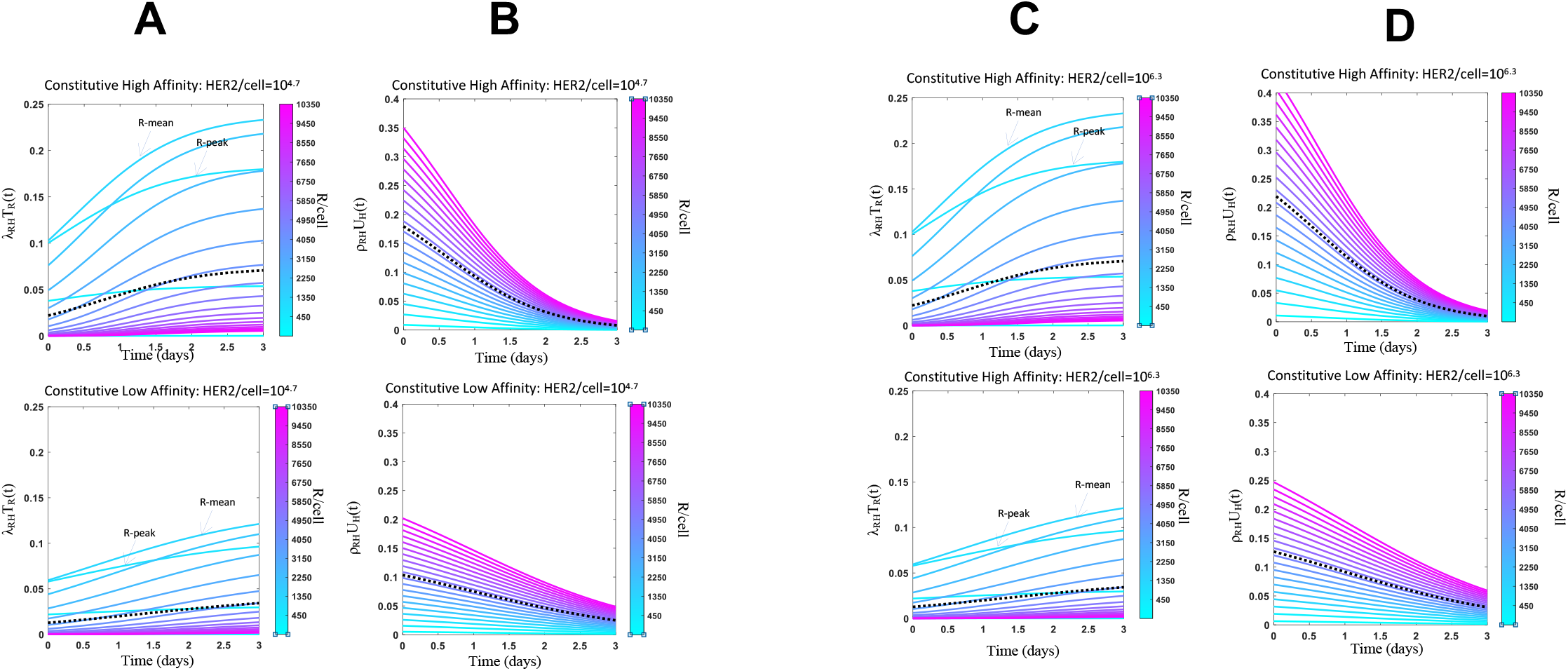
Constitutive CAR-T cell subpopulations expressing different CAR abundances show different cytotoxic and proliferative responses. **(A, C)** Shows lysis rates of target cells expressing different mean HER2 abundances (10^4.5^ vs 10^6.2^ molecules/cell) by subpopulations of CAR-T cells expressing different CAR abundances at different times post co-incubation. Results are shown for CARs with high (17.6 nM, top panel) and low (210.0 nM, bottom panel) affinity towards HER2. The dotted line shows the average of lysis rates across the CAR-T cell subpopulations. **(B, D)** Shows proliferation rates of CAR-T cells as they interact with target cells expressing different mean HER2 abundances (10^4.5^ vs 10^6.2^ molecules/cell) for the same assays in A and C.

**Figure S3.**
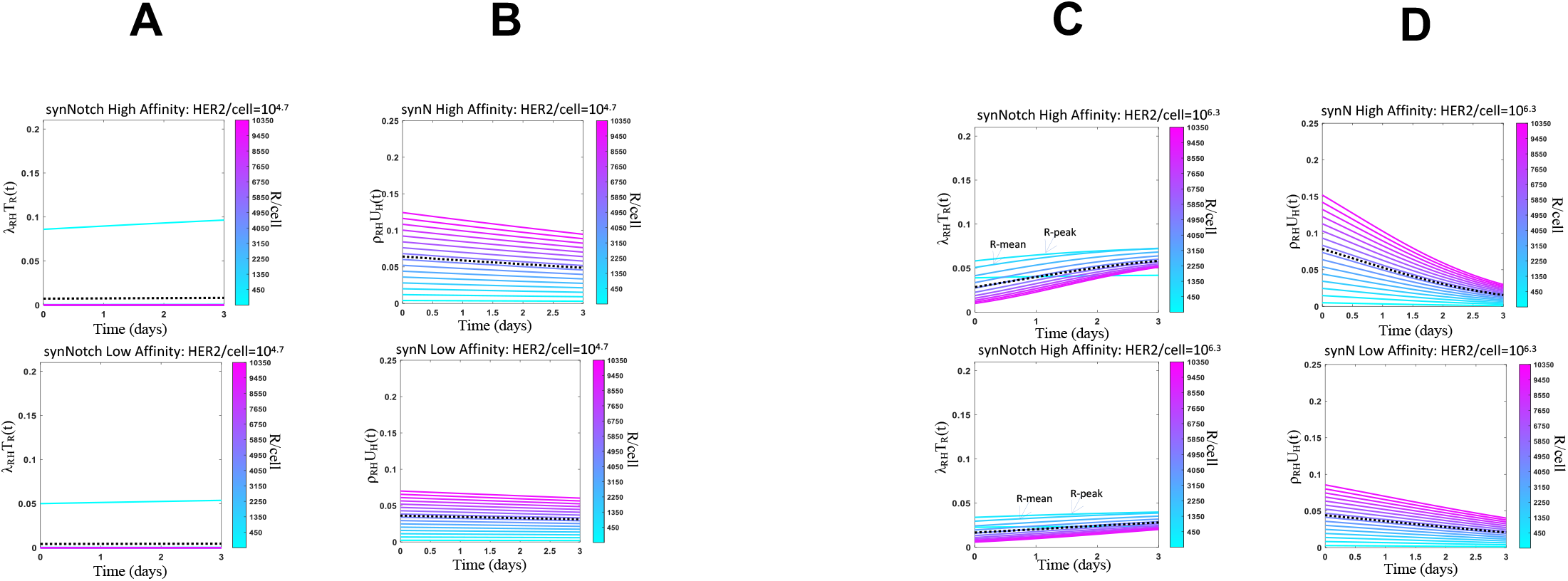
PASCAR modeling of cytotoxic and proliferation responses of synNotch CAR-T cell against target cells. **(A, C)** Shows lysis rates of target cells expressing different mean HER2 abundances (10^4.5^ vs 10^6.2^ molecules/cell) by subpopulations of synNotch CAR-T cells expressing different CAR abundances at different times post co-incubation. Results are shown for CARs with high (17.6 nM, top panel) and low (210.0 nM, bottom panel) affinity towards HER2. The dotted line shows the average of lysis rates across the synNotch CAR-T cell subpopulations. **(B, D)** Shows proliferation rates of synNotch CAR-T cells as they interact with target cells expressing different mean HER2 abundances (10^4.5^ vs 10^6.2^ molecules/cell) for the same assays in A and C.

**Figure S4.**
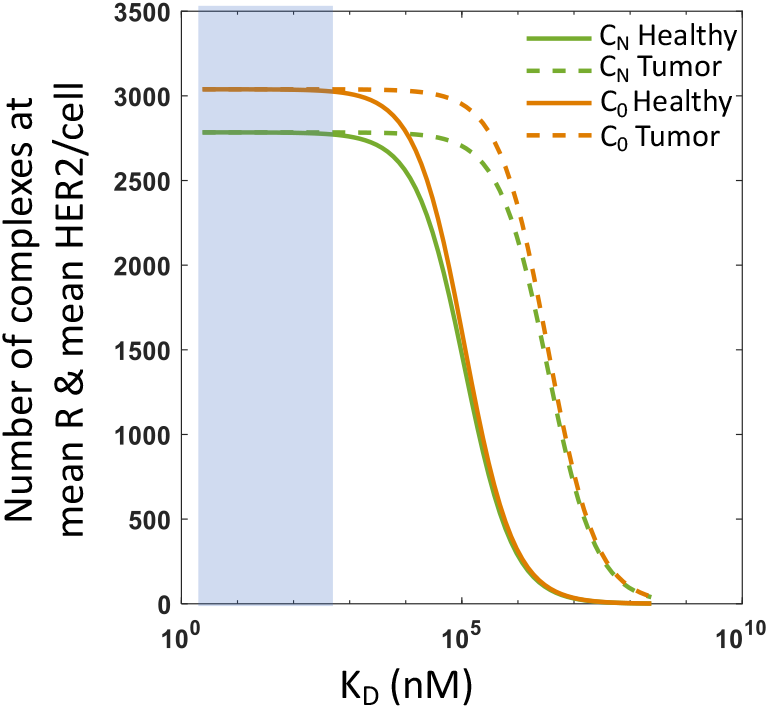
Variation of the steady state abundances of C_0_ and C_N_ with K_D_. Shows the variation of *C*_0_ and *C*_N_ with K_D_ where the abundances are computed at mean CAR= 3038 and mean HER2 abudances for healthy (=10^4.5^) and tumor (=10^6.2^) cells. The k_off_ and k_p_ are set to 9x10^−5^ s^-1^ and 0.0072. *C*_0_ and *C*_N_ do not change appreciably when CAR-T cells interact with healthy or tumor cells for smaller K_D_ values (< 100 nM).

**Figure S5.**
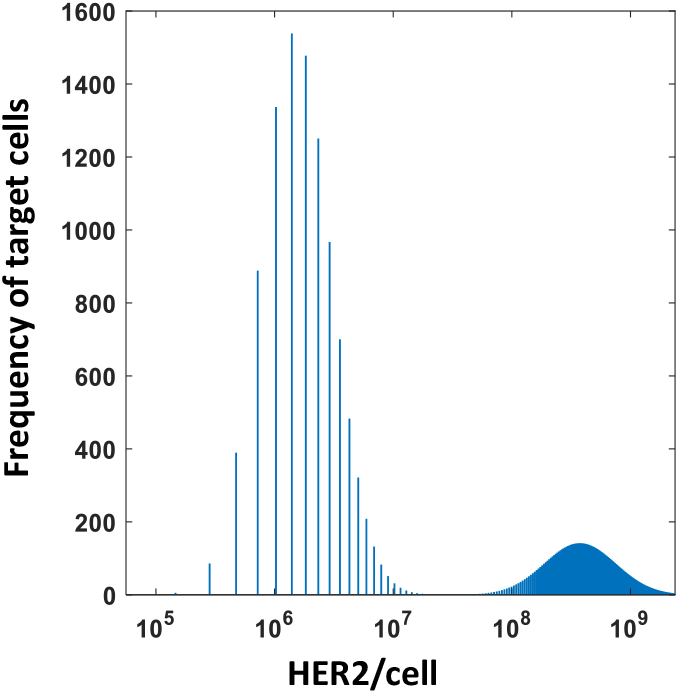
Simulated initial distribution of HER2 densities per target cell in a population of healthy and tumor cells. Shows normalized distribution of HER to in a mixture of healthy (10^4.5^ molecules/cell on average) and tumor cells (10^6.2^ molecules/cell on average) at t=0. The distribution is a superposition of two log normal distributions with means at 10^4.5^ molecules/cell and 10^6.2^ molecules/cell on average and standard deviation of 0.3.

